# Main compounds in sweat, such as squalene and fatty acids, should be promising pheromone components for humans

**DOI:** 10.1101/2024.08.15.608030

**Authors:** Yao-Hua Zhang, Yu-Feng Du, Jian-Xu Zhang

## Abstract

Pheromones are chemicals released outside the body by organisms to transmit information between individuals of the same species, thereby regulating behavior and physiology. Many biological and psychological studies have shown that human sweat does indeed contain chemical information related to gender, sex, emotion, etc., but there is no convincing answer to its chemical components. We conducted a GC-MS analysis of the chemical composition of adult and child sweat of Han Chinese, and found that the main components were squalene and fatty acids, and there were sex differences in adults, but not in children. Based on our long-term research on the coding rules of pheromones in a variety of animals, especially rodents, as well as men having more sweat glands and sebaceous glands, we speculate that squalene and many common fatty acids are likely to encode olfactory information such as sex and emotion in one component or a mix of components or in a dose-dependent manner. We also discussed that the main olfactory system and olfactory learning in social interactions should play an important role in human pheromone perception.

## INTROUCTION

Pheromones (intraspecific chemo-signals) are chemicals or a blend of several chemicals secreted to the outside of an individual and received by a second individual of the same species. These signals elicit a specific behavioral or physiological reaction (Karlson and Lüsche 1959; Wyatt 2014). The first pheromone (bombykol) was discovered to be a single component isolated from female silkworm moths in 1959 (Butenandt et al. 1959). This discovery led to the understanding that a pheromone can be composed of multiple chemical components and that pheromones can have synergistic effects (Silverstein et al. 1966,1967). So far, pheromones have been shown to exist in all animal groups, from protozoa to primates. Even birds, previously thought to rely solely on visual and auditory cues, have recently been confirmed to have pheromones (Novotny et al. 1999; Hagelin et al. 2003; Novotny 2003; Zhang et al. 2010; El-Sayed 2014; Zhang et al. 2014; Wyatt 2014). The animal group with the best studied pheromones are insects, and their pheromones are widely used in pest control (El-Sayed 2014; Wyatt 2014). The study of insect pheromones and their communication has laid the foundation of the main theories and research techniques in chemical ecology and animal pheromone communication (Butenandt et al. 1959; Silverstein et al. 1966, 1967; information exchange in organism kingdom, therefore, the pheromone components of these signals should have a common origin and phylogenetic similarity across species (Stoka 1999; Zhang et al. 2003, 2005; Zhang et al. 2008a; Xiao et al. 2009, 2010; Sandra et al. 2011; El-Sayed 2014; Wyatt 2014).

### Mammalian pheromones and their potential sources

Pheromones are chemical compounds animals release externally, and are mainly produced by exogenic glands, excreta, body surface compounds, etc (Wyatt 2014). Pheromones can be divided into a variety of functional types, for example, sex pheromones, alarm pheromones, aggregation pheromones, kin pheromones, social rank pheromones, territory pheromones, priming pheromone and so on (Wyatt 2014). In fact, a similar chemical signal exists between species, called interspecific chemosignals (allomone and kairomone) (Wyatt 2014). Airborne pheromones, the primary and most classic pheromone signals, are small and volatile to act at a distance, like sex attractants and alarm pheromones (Brennan and Zufall 2006; Brennan et al. 2010; Wyatt 2014). Additionally, non-volatile proteins and peptides can also act as pheromones, especially in aquatic animals and mammals (Brennan and Zufall 2006; Brennan et al. 2010; Wyatt 2014). In the case of mice and rats, many known volatile pheromone components from urine and preputial glands are the main components of these two sources of odor (Novotny et al. 1999; Novotny 2003; Zhang et al. 2008a,b; Zhang and Zhang 2014; Liu et al. 2019).

Extensive research suggests that mammalian pheromones come from exocrine glands like specialized skin glands, preputial glands, and lacrimal glands, as well as from metabolites like those found in urine (Johnston 1993; Novotny 2003; Kimoto et al. 2005; Zhang and Zhang 2014). In humans, these exocrine glands include sebaceous glands, eccrine sweat glands, apocrine sweat glands, lacrimal glands, salivary glands, mammary glands and so on, where the first two are much more widely distributed than others, secrete sebum- or water-based substances onto the skin’s surface (Kurosumi et al. 1984). Most studies on body odor in Caucasians and Africans focus on their strong-smelling axillary apocrine sweat glands, while Asians often have much smaller apocrine glands, resulting in only a faint acidic odor in the armpits (Martin et al. 2010). Therefore, if human pheromones exist, these two abundant exocrine glands (sebaceous glands and eccrine sweat glands) could be potential sources of common body odor and pheromones, especially important for Asians with the weakest axillary odor (Karlson and Lüscher 1959; Martin et al. 2010; Wyatt 2014).

### Sebaceous gland secretions may have pheromone components

Sebaceous glands are holocrine glands located in the dermis; they open into hair follicles and secrete a viscous, lipid-rich fluid (oily substance/sebum), which lubricates your hair and skin (Inoue et al. 2005; Baker 2019). Sebaceous glands are present over most of the body surface, particularly concentrated on the scalp, forehead, face, and anogenital area (Porter 2001; Sugawara et al. 2019). The human skin has an average of 2 million sebaceous glands, with a density reaching 400 to 900 glands per cm² on the face, these glands synthesize and secrete sebum, a key component of the hydrolipidic film protecting our skin (Baker 2019). Sebaceous glands and their pores are more abundant and larger in young man compared to young women. and shrink with age, similar to the specialized sebaceous glands in rodents like 2019; Novotny 2003). Human sebaceous glands, like the flank glands of golden hamsters and preputial glands of mice, are primarily controlled by androgens; they become active again at puberty, and sebum production is significantly higher in males compared to age-matched females (Pochi et al. 1979; Liu et al. 2010; Zhang et al. 2008a; Abdallah et al. 2017; Sugawara et al. 2019). Sebaceous gland secretions may act as pheromone in addition to their antibacterial and antifungal properties (Baker 2019).

Skin lipids, mainly produced by sebaceous glands, exhibit a wide range of both saturated and unsaturated fatty chains, creating a unique odor profile for each individual; these lipids differ from those produced by internal tissues, with skin producing odd chains, branched chains, and free acids, etc. (Nicolaides 1974). By the time sebum reaches the skin surface, bacteria have largely hydrolyzed the triglycerides, resulting in approximately one-third of the surface fat consisting of free fatty acids, which are potential components of human pheromones (Wertz 2007; Abdallah et al. 2017). Human sebum is reported to be a complex mixture of lipids—triglyceride fats (57.5 %), wax esters (26 %), squalene (12 %), cholesterol esters (3 %) and cholesterol (1.5 %) (de Lacy Costello et al. 2014; Baker 2019). Given that previous research on other mammals, especially rodents, has identified the main odor components as pheromones detectable by GC-MS (Novotny et al. 1999; Novotny 2003; Zhang et al. 2008 a,b; Liu et al. 2010, 2019). Squalene and fatty acids have been also identified as pheromone components in animals, we thus speculated that squalene and fatty acids from human sebaceous glands may also function as human pheromones (Mason et al. 1989; Yoder et al. 1993,1999; Pageat 2002; Zhang et al. 2008b; El-Sayed 2014).

### Body sweat may convey sexual and emotional information

Eccrine sweat glands are distributed across the body except for hair follicles, hair arrector muscles and sebaceous glands, are most concentrated on the palms and soles, followed by the head, with significantly lower density on the trunk and the extremities, which produce and secrete watery substances through ducts to cool the skin and are responsible for most sweat production (Kurosumi et al. 1984; Baker 2019).In contrast, apocrine sweat glands are primarily found in the axillae (armpits) and perineal area, contributing less to overall sweat production (Kurosumi et al. 1984).

Humans have more eccrine sweat glands than any other mammal, with the highest concentration and largest glands on the face and anogenital areas, which respond to both emotional and thermal stimuli (Wertz 2007; Wilke et al. 2007; Kamberov et al.2015; Baker ·2019).

Although men and women have the same number of sweat glands, due to the influence of testosterone, men have a larger size and volume of each sweat gland than women and produce about five times as much sweat (Baker 2019; Kawahata 1960; Inoue et al. 2005; Ichinose-Kuwahara et al. 2010). Functional magnetic resonance imaging (fMRI) have shown that the right orbitofrontal cortex, right fusiform cortex, and right hypothalamus respond to airborne-human sexual sweat, suggesting that sweat carries sexual and emotional information (Zhou and Chen 2008). Human sweat contains various fatty acids, including tetradecanoic acids, hexadecenoic acid and oleic acid, which can attract mosquito (Bernier et al. 1999, 2000; Verhulst et al. 2013; Zhang et al. 2022). These fatty acids, along with those from sebaceous glands, may be potential candidates for human pheromone components (El-Sayed 2014; Pageat 2002).

### The known chemical composition of human pheromones is highly controversial

The first human pheromones to be demonstrated were involved in menstrual cycle synchronization, and subsequent research revealed that odorless compounds from the armpits of women in the late follicular phase of their menstrual cycles accelerated the preovulatory surge of luteinizing hormone in recipient women and shortened their menstrual cycles, but those collected later in the menstrual cycle (at ovulation) had the opposite effect, providing highly persuasive evidence for the existence of human pheromones (McClintock 1971; Stern and McClintock 1998). Extensive research has also shown that human sweat carries chemical signals related to emotions such as fear, disgust and stress, for example, individuals experiencing stress sweet more and produce a distinctive odor that can induce a stress response in others, suggesting that human sweat may contain fear pheromones (Hays 2003; Wysocki and Preti 2004; Albrecht et al. 2011; de Groot et al. 2012, 2015, 2020; de Groot and Smeets 2017; Lübke et al. 2017; Wunder et al. 2023).

Pheromones in humans may be present in bodily secretions such as urine, semen, vaginal secretions, breast milk, and potentially saliva and breath; however, most attention thus far has been directed toward axillary sweat, because the underarm apocrine sweat glands are generally regarded as scent glands involved in the production of pheromones due to their smelly secretions (Hays 2003; Verhaeghe et al. 2013; Wyatt 2015; Baker LB. 2019).

Axillary secretions originate from the highly dense eccrine and apocrine sweat glands, as well as sebaceous glands (Baker 2019). Apocrine sweat glands open into hair follicles and produce viscous, lipid-rich sweat containing proteins, sugars, and ammonia (Hodge et al. 2022). Apocrine sweat is odorless, but it acquires odor after interaction with cutaneous bacteria, which, for example, act on the 16-androstenes to produce odorous volatiles (Verhaeghe et al. 2003; Havlicek et al. 2010; Mogilnicka et al. 2020). However, the identified chemical components of human pheromones remain highly controversial (Wysocki and Preti 2004; Wyart et al. 2007; Havlicek et al. 2010; Petrulis 2013; Doty 2014; Wyatt 2015; Hare et al. 2017). For chemical studies of human pheromones, the main focus has been on two axillary sweat steroids: androstenone and androdienone; some experiments have shown these steroids to have an effect on people through the sense of smell, while others have not (Nixon et al. 1988; Bird and Gower 1982; Gower et al. 1985, 1994; Lundström et al. 2006; Zhou and Chen 2008; Verhaeghe et al. 2013; Zhou et al. 2014; Ferdenzi et al. 2016). Steroid hormone metabolites secreted into the environment may act as pheromones in fish and mammalian (Melrose et al. 1971; Stacey and Sorensen 2009). In fishes, soluble steroids that act as hormones or their metabolites have been widely proven to serve as sex pheromones, termed “hormonal pheromones” (Stacey and Sorensen, 2009). Similarly, in boars, saliva is the major source of the pheromone androstenone (roughly 0.030 ug/mL) that can elicit mating stance behavior in sows in heat (Melrose et al. 1971). However, strong evidence for chemical communication via steroids among humans is lacking (Wysocki and Preti 2004; Havlicek et al. 2010; Petrulis 2013; Wyatt 2015; Hare et al. 2017).

If human sex pheromones influence our sexual attractiveness or emotions, then they are mammals, rather than steroids (Wyatt 2015). In fact, subsequent chemical analysis until today has found many volatile chemicals, mainly low molecular weight acids, aldehydes, alkanols, etc., in armpit sweat, but no such steroids have been detected (Zeng et al. 1991,1992, 1996; Zhang et al. 2005; Penn et al. 2007). Notably, repeated chemical analyses have shown that human male axillary odors consist of C_6_ to C_11_ normal, branched, and unsaturated aliphatic acids, with (*E*)-3-methyl-2-hexenoic acid being the most abundant; therefore, fatty acids are among the important components or major components of human body odor (Zeng et al. 1991,1992, 1996). A SNP in OR51B2 associated with trans-3-methyl-2-hexenoic acid is also found (Li et al. 2022). Carboxylic acid, especially (*E*)-3-methyl-2-hexenoic acid could be considered potential chemical signals in humans for certain functions, worthy of further verification (El-Sayed 2014; Pageat 2002).

The sweat on the skin surface of most parts of the human body mainly consists of secretions from sweat glands and sebaceous glands (Inoue et al. 2005; Baker 2019). If human pheromones do exist, sweat would be an important source of them, particularly among Chinese people known for having the weakest armpit odor (Martin et al. 2010; Baker et al. 2019). This paper analyzes the main volatile components of Chinese sweat associated with gender and age, and discusses potential components of human pheromones (Zou and Yang 2022; Qu et al. 2014).

## Materials and Methods

### 1.1. Sweat Donors

We recruited 31 undergraduate and graduate students as sweat donors, including 12 men, 10 women (excluding menstruation), and 9 women (during menstruation), aged 20 to 28. All donors were in good health and participated in gym classes during the 2009-2010 period.

Additionally, 26 children (13 boys and 13 girls) aged 6 to 10 were included. They were sweating profusely while playing or engaging in moderate physical activity. With parental consent, sweat samples were collected by the parents, ensuring that the children did not apply makeup during the collection process.

### 1.2. Sweat Collection and Extraction of Its Components

Medical cotton swabs were soaked in dichloromethane (purity >99.5%) for 12 hours to clean them, then dried, sealed in glass vials, and weighed for use. During sweat sampling, we gently used the swabs to absorb sweat from each donor’s forehead, sealed the swabs in 2 ml glass vials, and immediately stored them in an ice box. In the laboratory, the vials were weighed to determine the sweat weight, which ranged from a few milligrams to several dozen milligrams.

To extract compounds from each sweat sample, we added dichloromethane to each vial at a ratio of 1 mg of sweat to 30 µl of dichloromethane (purity >99.5%). The solution was kept at 0°C for 12 hours. After removing the swab, the remaining solution was stored at -20°C until it was used for GC–MS analysis.

### 2.3 Optimizing Conditions for GC/MS Analyses

The analysis was performed using an Agilent Technologies Network 6890N GC System coupled with a 5973 Mass Selective Detector (NIST 2002 Library). Pilot samples were similar to those of preputial gland secretions from laboratory rats (*Rattus norvegicus*). Therefore, we used the GC-MS conditions previously employed for analyzing rat preputial gland secretions (Zhang et al. 2008b, Zhang and Zhang 2014) to analyze the sweat volatile compounds.

Specifically, the GC was equipped with an HP5MS glass capillary column (30 m long, i.d. 0.25 mm × 0.25 µm film). Helium was used as the carrier gas at a flow rate of 1.0 ml/min. The injector temperature was set to 280°C. The initial oven temperature was set to 50°C, increased by 5°C/min to 100°C, then ramped by 10°C/min to 280°C, and held at 280°C for 5 minutes. Electron impact ionization was performed at 70 eV. The transfer line temperature was maintained at 280°C. The scanning mass range was from 30 to 450 amu. A 2 µl sample was injected in splitless mode.

Tentative identifications were made by matching the mass spectra of GC peaks with those in the MS library (NIST 2002). Among the tentatively identified compounds, indole, *E*-*ß*-farnesene, tetradecanoic acid, hexadecanoic acid, octadecanoic acid, squalene, cholesterol, 2-heptanone, and 4-ethyl phenol were verified by matching the retention times and mass spectra with authentic analogs (all purities >95%, purchased from J&K Chemical Ltd, Beijing, China). Additionally, 4-heptanone and *E*,*E*-α-farnesene were verified by comparison with their counterparts in mouse urine and preputial glands (Zhang, Rao, et al. 2007).

We quantified the abundance of each compound by its GC area. The peak area of each compound was converted into a percentage of the summed peak areas from all targeted GC peaks of either urine or preputial glands, representing its relative abundance. These abundances and relative abundances of the scent constituents were used for quantitative comparisons between men and women or between boys and girls. If a GC peak was too small to display the diagnostic MS ions of the corresponding compound, its GC area was recorded as zero. This approach was also applied to GC peaks below the detectable limit.

## Results

### 1. Volatile components of adult sweat and their sex differences

A total of 46 components were detected in men and women sweat. In man sweat, carboxylic acids accounted for 21.43%, squalene for 84.28%, and other components (aldehydes, alkanes, terpenes, esters, etc.) for 5.09%. In woman sweat, carboxylic acids, squalene, and other components accounted for 20.60%, 69.85%, and 9.55%, respectively. In menstruating women, these compounds accounted for 20.60%, 69.85%, and 9.55%, respectively. Squalene was the most abundant compound detected in all adult sweat samples. Among the 20 carboxylic acids detected, tetradecanoic acid, pentadecanoic acid, hexadecanoic acid, and palmitoleic acid were the most abundant.(Fig 1; Table 1)

**Figure 1.**
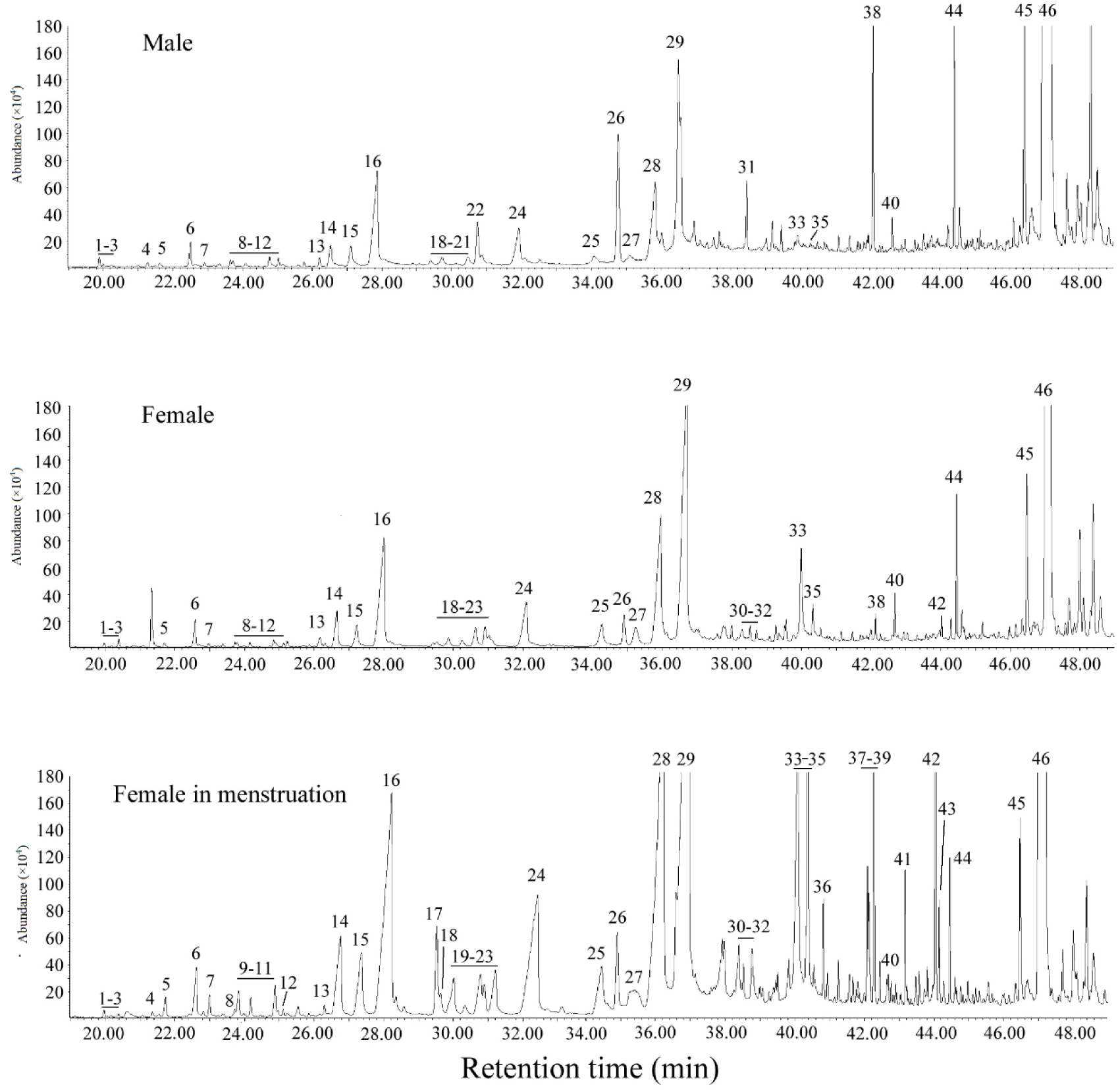
Representative GC profile of dichloromethane extract from man and woman sweat. GC conditions are described in materials and methods section. Numbered GC peaks correspond to compounds in Table 1.

**Figure 2:**
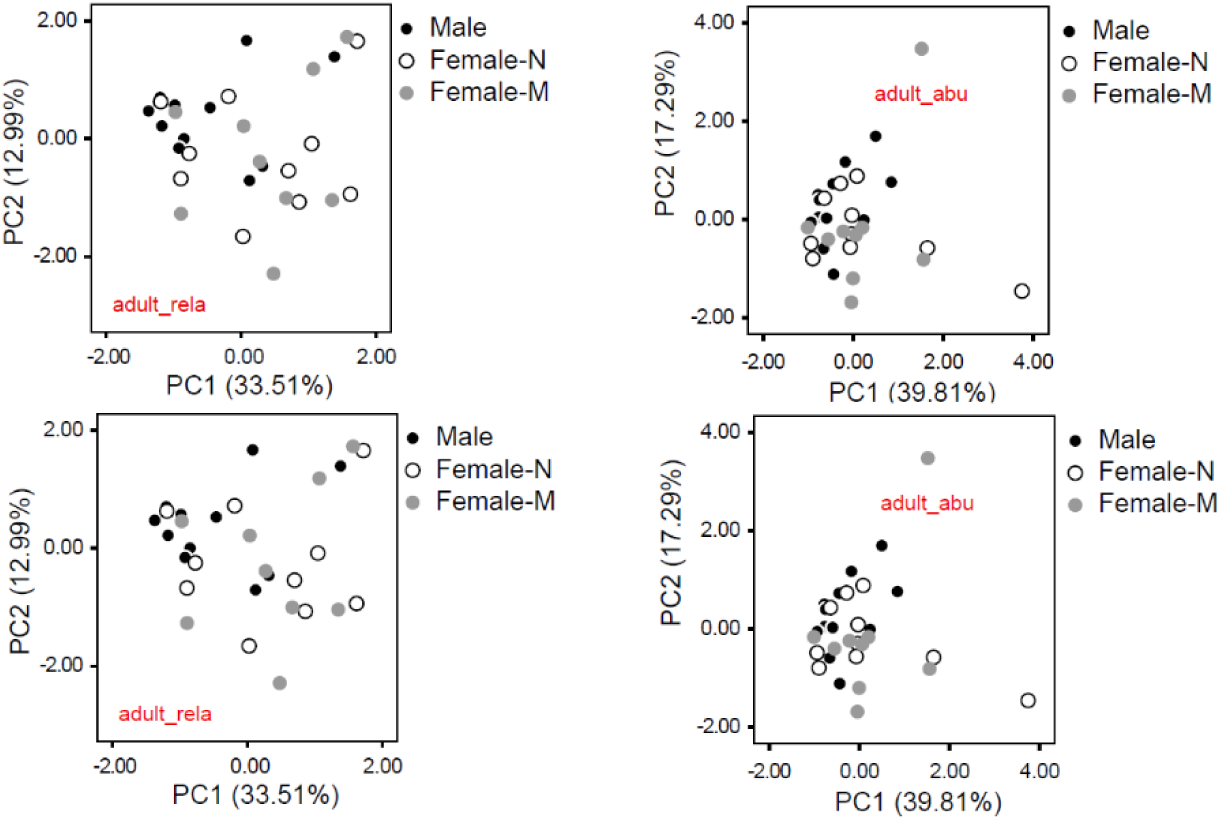
Principal component plots of men (black dots) and women (excluding menstruation) (blank dots) and women with menstruation (gray dots). Each symbol represents one person (Top left panel: Relative abundance of all compounds; Top right panel: Abundance of all compounds; Bottom left: Relative abundance of only carboxylic acids; Bottom right: relative abundance of only carboxylic acids)

**Table 1.**
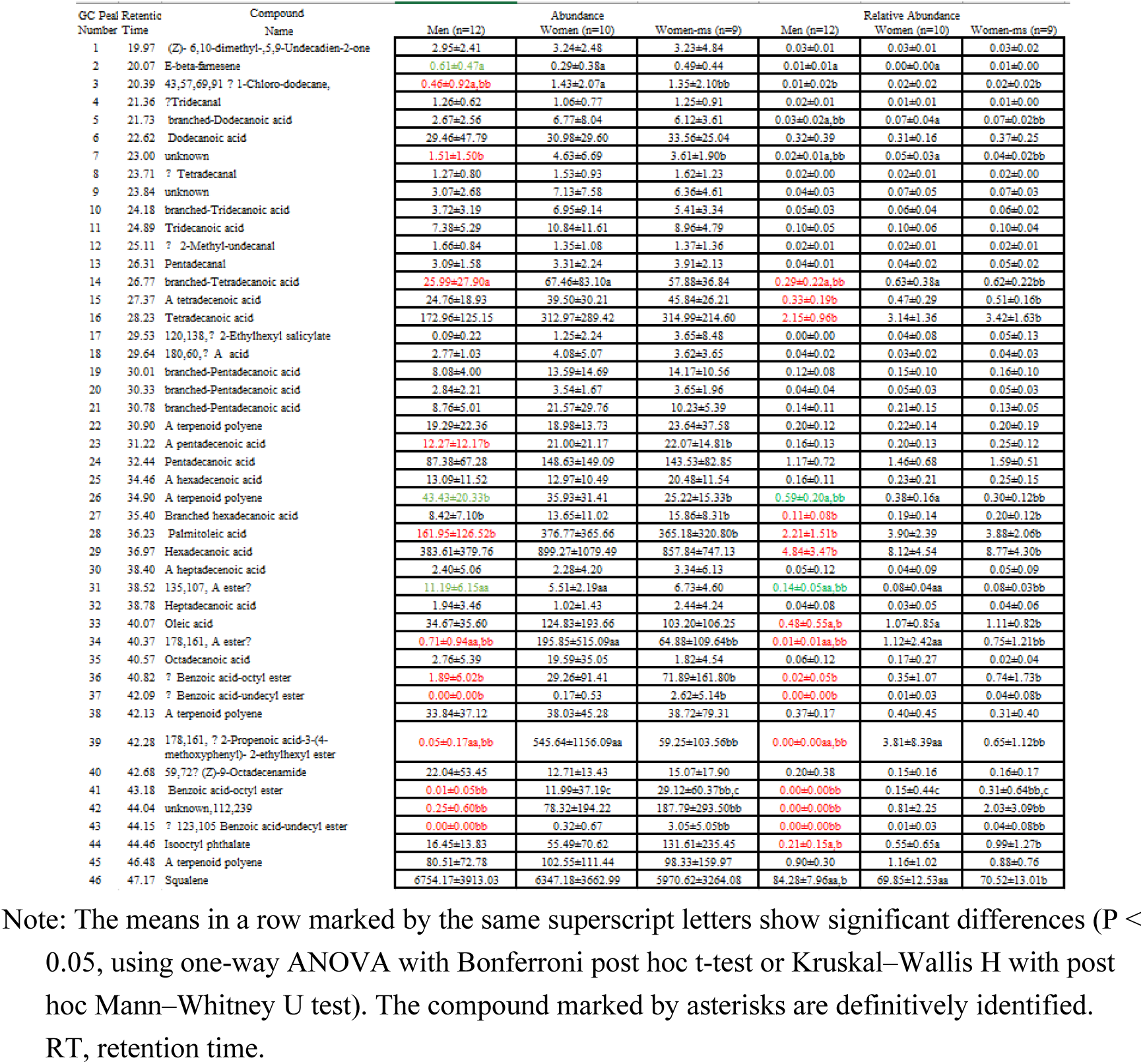
Comparison of abundance or relative abundance of sweat volatiles between men and women (Mean±standard deviation, n=12 for men, 10 for women (excluding menstruation), and 9 for women with menstruation).

Both the absolute and relative abundances of seven carboxylic acids, including tetradecanoic acid, a tetradecenoic acid, a branched tetradecanoic acid, hexadecanoic acid, a hexadecenoic acid, a branched hexadecanoic acid, and oleic acid, were higher abundances of seven other components, including esters, terpenes, and amides, were higher in women. Conversely, squalene, along with *E*-β-farnesene, a terpenoid polyene and an ester, had greater abundance and/or relative abundant in men than in women.

In the principal component analysis, based on both absolute and relative abundances of all detected volatile compounds or only carboxylic acids, there was almost complete overlap between men and women (Fig2)

### 2. Volatile components of children’s sweat and their sex differences

A total of 40 compounds were detected in boys’ and girls’ sweat, including 22 carboxylic acids, squalene, and others (aldehydes, alkanes, terpenes, etc.). The most abundant acids were tetradecanoic acid, pentadecanoic acid, hexadecanoic acid, palmitoleic acid, oleic acid, and octadecanoic acid. In boys’ sweat, carboxylic acids accounted for 39.10%, squalene for 55.43%, and other components for 7.47%. In girls’ sweat, carboxylic acids, squalene, and other components accounted for 42.43%, 47.04%, and 10.53%, respectively.( Fig 3; Table 2)

**Figure 3.**
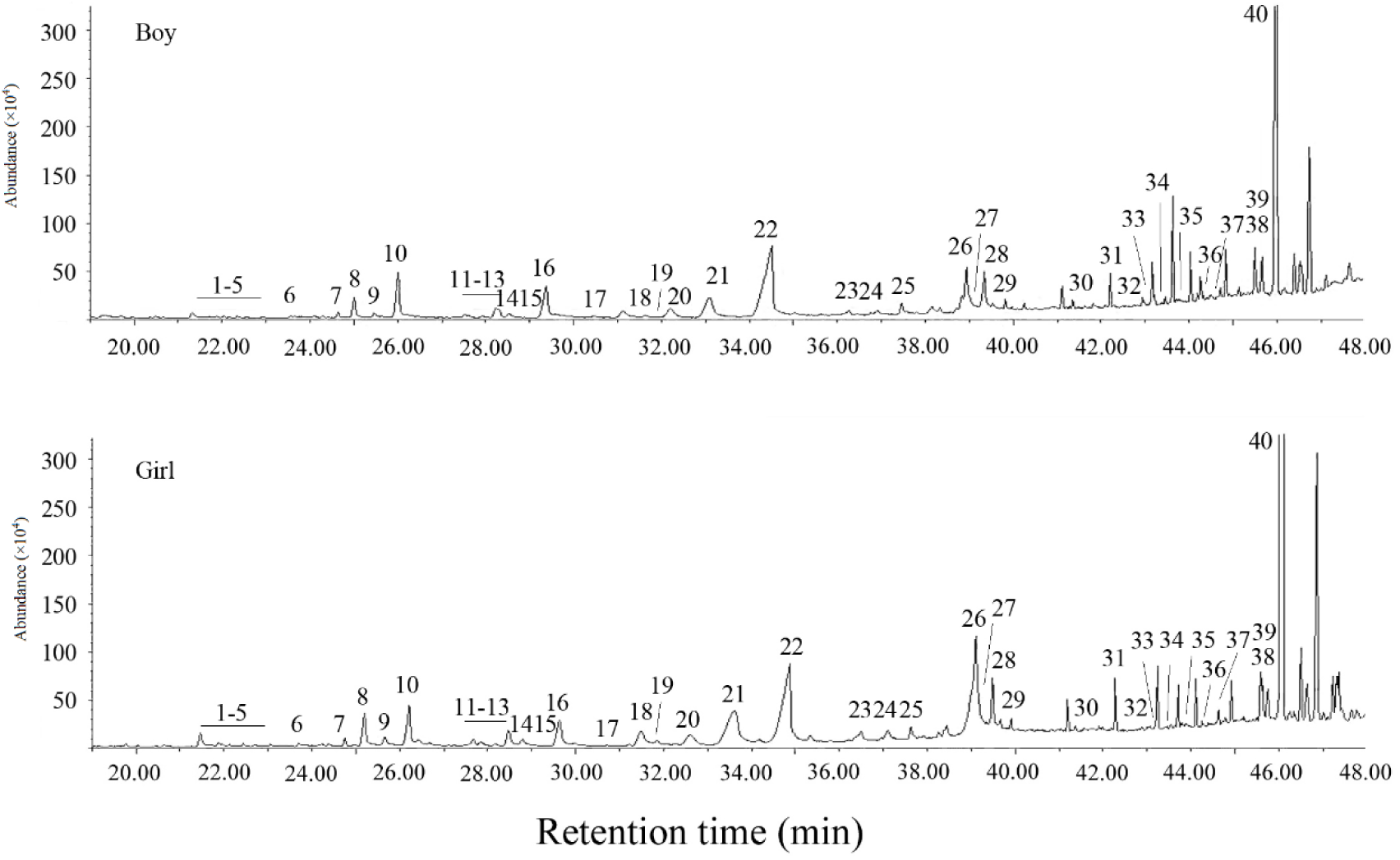
Representative GC profile of dichloromethane extract from boys’ and girls’ sweat. GC conditions are described in materials and methods section. Numbered GC peaks correspond to compounds in Tables 2.

**Table 2.** Comparison of abundance or relative abundance of sweat volatiles between boys and girls (Mean±standard deviation, n=13 for either boys or girls).

| GC Peak Number | Retention Time | Compound Name | Abundance |  | Relative Abundance |  |
| --- | --- | --- | --- | --- | --- | --- |
|  |  |  | Boys (n=13) | Girls (n=13) | Boys (n=13) | Girls (n=13) |
| 1 | 21.24 | Dodecanoic acid | 3.36±3.70 | 3.29±3.61 | 0.65±0.49 | 0.60±0.20 |
| 2 | 21.66 | A branched-Tridecanoic acid | 0.13±0.47 | 0.13±0.33 | 0.01±0.05 | 0.01±0.03 |
| 3 | 22.30 | Tridecanal | 0.65±0.80 | 0.55±0.32 | 0.13±0.11 | 0.14±0.10 |
| 4 | 22.46 | A branched-Tridecanoic acid | 0.20±0.41 | 0.12±0.23 | 0.02±0.05 | 0.02±0.03 |
| 5 | 22.82 | A branched-Tridecanoic acid | 0.25±0.43 | 0.16±0.31 | 0.03±0.04 | 0.02±0.04 |
| 6 | 23.47 | Tridecanoic acid | 0.67±1.23 | 0.29±0.44 | 0.06±0.12 | 0.03±0.05 |
| 7 | 24.59 | Tetradecanal | 1.97±1.90 | 1.86±1.17 | 0.43±0.28 | 0.49±0.32 |
| 8 | 24.90 | A branched-Tetradecanoic acid | 4.62±7.53 | 3.86±5.94 | 0.74±0.76 | 0.55±0.52 |
| 9 | 25.36 | A Tetradecenoic acid | 1.53±1.99 | 0.86±1.53 | 0.14±0.20 | 0.11±0.18 |
| 10 | 25.86 | Tetradecanoic acid | 19.47±17.58 | 14.04±11.64 | 3.19±1.43 | 2.74±0.99 |
| 11 | 27.45 | Hexadecanal and branched-Pentadecanoic acid | 2.54±2.08 | 1.89±1.33 | 0.47±0.22 | 0.42±0.18 |
| 12 | 27.78 | A branched-pentadecanoic acid | 0.87±0.84 | 0.64±0.87 | 0.14±0.10 | 0.08±0.06 |
| 13 | 28.07 | A branched-pentadecanoic acid | 2.56±2.86 | 2.14±3.26 | 0.32±0.25 | 0.33±0.32 |
| 14 | 28.21 | A terpenoid polyene | 8.05±14.04 | 5.89±13.19 | 0.91±0.58 | 0.71±0.59 |
| 15 | 28.42 | A pentadecenoic acid | 1.50±2.19 | 0.67±1.10 | 0.20±0.23 | 0.12±0.15 |
| 16 | 29.14 | Pentadecanoic acid | 15.17±15.95 | 9.03±8.32 | 2.26±1.31 | 1.66±0.87 |
| 17 | 29.86 | Hexadecanol | 18.84±60.24 | 36.93±101.46 | 2.07±6.39 | 2.61±5.57 |
| 18 | 30.91 | A Hexadecenoic acid | 2.26±3.89 | 2.07±4.50 | 0.36±0.52 | 0.23±0.45 |
| 19 | 31.47 | A terpenoid polyene | 1.17±2.31 | 0.56±0.91 | 0.10±0.16 | 0.08±0.15 |
| 20 | 31.94 | A branched-hexadecanoic acid | 4.21±4.34 | 3.22±4.62 | 0.71±0.64 | 0.51±0.44 |
| 21 | 32.82 | Palmitoleic acid | 15.93±15.09 | 11.91±16.74 | 2.46±1.96 | 1.87±1.66 |
| 22 | 34.29 | Hexadecanoic acid | 108.23±102.36 | 97.14±94.06 | 20.48±11.55 | 20.86±11.23 |
| 23 | 36.16 | A branched-heptadecanoic acid | 1.73±2.19 | 1.36±1.97 | 0.26±0.27 | 0.23±0.24 |
| 24 | 36.81 | A heptadecenoic acid | 1.42±2.09 | 0.73±1.80 | 0.22±0.29 | 0.08±0.18 |
| 25 | 37.36 | Heptadecanoic acid | 3.37±3.44 | 3.85±4.84 | 0.64±0.53 | 0.72±0.47 |
| 26 | 38.92 | Oleic acid | 21.12±21.14 | 22.81±30.34 | 3.71±3.22 | 4.41±3.15 |
| 30 | 39.24 | 2-Ethylhexyl-p-methoxycinnamate (178, 161) | 1.34±4.84 | 14.08±39.39 | 0.29±1.03 | 2.20±5.16 |
| 27 | 39.48 | Octadecanoic acid | 13.44±16.24 | 20.08±45.13 | 2.08±1.26 | 2.82±2.34 |
| 31 | 39.75 | Benzoic acid-octyl ester (123, 105) | 0.00±0.00 | 5.99±21.36 | 0.00±0.00 | 0.82±2.70 |
| 32 | 41.32 | 2-Ethylhexyl-4-methoxycinnamate (178, 161) | 1.71±3.02 | 13.82±35.61 | 0.24±0.50 | 2.54±5.44 |
| 33 | 42.27 | Benzoic acid-octyl ester (123, 105) | 0.15±0.55 | 3.38±10.25 | 0.02±0.06 | 0.45±1.29 |
| 34 | 42.63 | Benzoic acid-octyl ester (123, 105) | 0.00±0.00 | 0.10±0.36 | 0.00±0.00 | 0.12±0.44 |
| 35 | 42.93 | Hexanoic acid-2-ethyl-hexadecyl ester (145) | 0.28±0.69 | 11.42±41.17 | 0.03±0.09 | 0.53±1.92 |
| 36 | 43.27 | Benzoic acid-octyl ester (123, 105) | 0.00±0.00 | 0.08±0.30 | 0.00±0.00 | 0.10±0.37 |
| 37 | 43.81 | A ester (145) | 0.00±0.00 | 0.18±0.67 | 0.00±0.00 | 0.02±0.08 |
| 38 | 44.18 | Benzoic acid-octyl ester (123, 105) | 0.00±0.00 | 4.32±12.74 | 0.00±0.00 | 0.55±1.61 |
| 39 | 45.44 | A terpenoid polyene | 20.33±32.47 | 15.84±21.39 | 3.21±2.56 | 2.91±2.31 |
| 40 | 45.91 | Squalene | 399.76±451.49 | 296.77±317.02 | 53.43±16.59 | 47.04±19.81 |

Note: The means in a row marked by the same superscript letters show significant differences ( $P < 0.05$ , using t-test or Mann–Whitney U test). RT, retention time.

The detected volatile compounds did not differ between boys and girls in either absolute or relative abundance. (Table 2)

In the principal component analysis, based on both absolute and relative abundances of all detected volatile compounds or only carboxylic acids, there was almost complete overlap between boys and girls.(Fig. 4)

**Figure 4:**
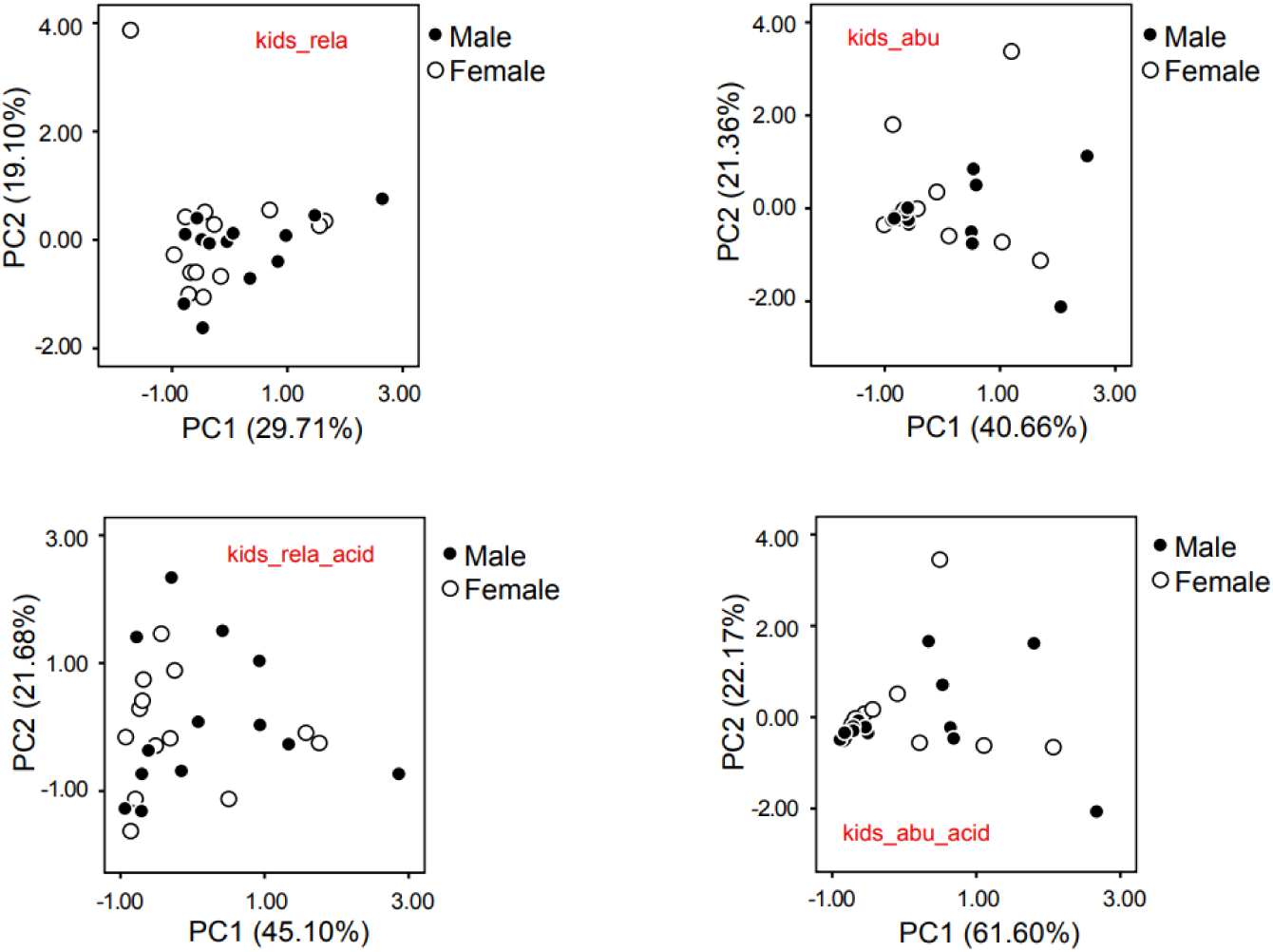
Principal component plots of males (boys) (black dots) and females (girls) (blank dots) and women with menstruation (gray dots). Each symbol represents one person (Top left panel: Relative abundance of all compounds; Top right panel: Abundance of all compounds; Bottom left: Relative abundance of only carboxylic acids; Bottom right: relative abundance of only carboxylic acids)

## DISCUSSION

Here, we demonstrated that squalene is the most abundant and male-enhanced component in Chinese sweat detected by GC-MS, though there was no separation between men and women in the principal component analysis, similar to its presence in rat preputial gland secretions measured under the same GC-MS conditions, (Zhang et al. 2008b; Zhang and Zhang 2014).We also found that some common aliphatic acids (saturated and unsaturated) are among the most abundant components in sweat, as previously reported (Bernier et al. 1999, 2000; de Lacy Costello et al. 2014). Despite no separation between men and women in principle component analysis

### 1. Squalene as the Most Abundant and Masculine Component of Sweat May Be a Male Pheromone Component

Squalene has been previously identified as the main component of skin surface polyunsaturated lipids, acting as an emollient and antitumor compound (Bernier et al. 1999, 2000; Amarowicz 2009; Huang et al. 2009). It has been shown to act as a pheromone or interspecific chemical signal in snakes, insects, rats, and other species (Mason et al. 1989; Yoder et al. 1993, 1999; Zhang et al. 2008b; López and Martín 2009; El-Sayed 2014; Yang et al. 2017). This suggested that squalene, despite being a semi-volatile compound, is strong enough to be detected by animals at a distance and can therefore be used as olfactory signals in animals, and might serve as a component of a pheromone in humans.

Mammalian pheromones are best identified in the urine and preputial glands of (S)-2-sec-butyl-4,5-dihydrothiazole, *R*,*R*-3,4-dehydro-exo-brevicomin, and 6-hydroxy-6-methyl-3-heptanone in male mouse urine, and *E*-β-farnesene, *E*,*E*-α-farnesene, hexadecanol, and hexadecyl acetate in male mouse preputial glands, as well as 2-heptanone in male rat urine (Novotny et al. 1999a, b; Zhang et al. 2007; 2008a, b; Zhang and Zhang 2014). The correlation between the composition of volatile components of odors released outside the body and biological properties such as gender, emotions, and physiological/social states is essential for identifying pheromones (Novotny et al. 1999b, 2003; Wyatt 2014). For example, sex pheromone components must be male/female-unique (digital code) or male/female-enhanced (analogue code) volatile compounds (Novotny et al. 1999a, b; Zhang et al. 2008a, b, c; Liu et al. 2010). Therefore, when considering human pheromones, the main components in body odor should be prioritized.

Although we only showed that the relative abundances of sweat squalene were significantly higher in males than in females, men should produce more sebum and excrete more sweat due to the larger and more numerous sebaceous glands and sweat glands than women did females, therefore men must release much more squalene than women to be a potential male pheromone in humans (Abdallah et al. 2017; Baker 2019; Sugawara et al. 2019).

We also observed that the aforementioned sexual differences disappeared in children, indicating that the composition of volatile chemicals in human sweat is related to sexual development, differentiation, and maturation, and is controlled by androgens (Kunkeler and Hodgins1994; Baker 2009). This further substantiates squalene’s identity as a male pheromone in humans (Novotny 2003; Zhang et al. 2008a, b; Zhang et al. 2019b). In fact, sebaceous and sweat glands have more androgen receptors in males than in females (Kunkeler and Hodgins 1994; Abdallah et al. 2017).

There are exceptions among mammals that do not rely on olfaction for sex recognition. For instance, the Asian house rat (*Rattus tanezumi*) has weak sex-differential volatile compounds in its urine and preputial glands, making it unable to attract the opposite sex through olfaction, however, ultrasound exhibits distinct sex dimorphism in this species (Wang et al. 2023). Nonetheless, human sweat should contain human sexual chemosensory cues (Zhou and Chen 2008).

### 2. Are aliphatic acids in sweat potential human sex pheromones?

These common fatty acids also possess the chemical properties of pheromones, which are widely used as pheromones in various insects (El-Sayed 2014; Pageat 2002).

In mammals, a mixture of fatty acids such as linoleic, oleic, and palmitic acids or their derivatives can act as appeasing pheromones to reduce stress, anxiety, and aggressiveness in species such as pigs, dogs, and horses (El-Sayed 2014; Pageat 2002; Tod et al. 2005; Yonezawa et al. 2009; Berger et al. 2013; Taylor et al. 2020; Kiyokawa et al. 2023). These fatty acids also function as priming pheromones in goats (Sugiyama et al. 1981; Hillbrick et al. 1995). A blend of fatty acids has been shown to bind with various carrier proteins in pig nasal mucus, confirming their potential role in the olfactory and vomeronasal systems (Pageat 2002; Yonezawa et al. 2009; Sankarganesh et al. 2022).

In Mongolian gerbils, phenylacetic acid from the mid-ventral gland secretion is a major male pheromone (Thiessen et al. 1974; Liu et al. 2010). In golden hamsters, tetradecanoic acid and hexadecenoic acid secreted from the flank glands act as female pheromones to attract males and reduce male-male aggression (Zhang et al. 2008c; Liu et al. 2010). In rhesus monkeys, vaginal short-chain fatty acids (e.g., acetic acid, propionic acid, iso-butyric acid, butyric acid, isovaleric acid, and isocaproic acid) can stimulate male sexual behavior and are considered sex pheromones (Curtis et al. 1971; Michael et al. 1971; Bonsall et al. 1980).

In humans, a series of fatty acids, including C13, C14, C16, C18, oleic acid, linoleic acid, and linolenic acid, are present in colostrum and areolar (Montgomery’s) gland secretions of women, as well as on human skin, which may act as Human Appeasing Pheromones (HAP) to influence psychological feelings through olfaction (Bernier et al. 1999, 2000; Schaal et al. 2008; Doucet et al. 2009; Pageat et al. 2014; Piccinni et al. 2018). Even in birds, a mixture of fatty acids or their derivatives from uropygial gland secretions can function as appeasing pheromones (Pageat 2010). This suggests that fatty acids possess the chemical properties necessary to be components of pheromones in both animals and humans.

On the other side, fatty acids in human body sweat have been proven to act as interspecific chemosignals. Human sweat fatty acids with fewer than 20 carbon atoms, such as tetradecanoic acid, hexadecenoic acid, and oleic acid, can function as interspecific chemosignals to attract mosquitoes (Bernier et al. 1999, 2000; Smallegange et al. 2009; Verhulst et al. 2013). People carrying the HLA gene Cw*07 have high levels of lactic acid, 2-methyl-butanoic acid, and tetradecanoic acid, making them highly attractive to mosquitoes (Verhulst et al. 2013). Dogs have a powerful sense of smell and can distinguish between individuals, health states (e.g., diabetes, cancer, epilepsy), or emotional states by smelling people’s sweat, which contains a complex mixture of volatile organic compounds such as fatty acids, acting as an odor fingerprint (Shirasu and Touhara 2011; Angle et al. 2016; Wilson et al. 2022). In fact, intraspecific and interspecific chemical signals can share some chemical components, or an odor component can participate in both intraspecific chemical communication and interspecific interactions through the sense of smell (Zhang et al. 2003, 2005, 2007; El-Sayed 2014; Wyatt 2014).

Since fatty acids in human sweat can be used as interspecific chemical signals for mosquitoes, dogs, and other species, they could also serve as intraspecific chemical signals (e.g., pheromone components) to communicate gender identity, physiological, and emotional states. Humans are sensitive to 3-methyl-2-hexenoic acid, a major component of human axillary odor, which might be involved in male axillary extracts affecting the pulsatile secretion of luteinizing hormone and mood in female recipients (Wysocki et al. 1993; Pierce et al. 1995; Preti et al. 2003).

In the current study, the relative or absolute abundance of these fatty acids in the same volume of sweat was greater in women than in men, implying potential female pheromone components in humans. This difference was not observed in children, suggesting that sexual development and differentiation might enhance these major fatty acids in female sweat. But, men’s sweat glands and sebaceous glands are much larger and more numerous than women’s, so men release more sweat and total fatty acids than women, therefore, these major fatty acids probably act as male pheromones at high doses in humans (Abdallah et al. 2017; Baker 2019; Sugawara et al. 2019).

This phenomenon is similar to what was found in mice, where urine-borne hexadecanol and hexadecyl acetate secreted from the preputial glands are common in both males and females. However, males produce these compounds in greater abundance, attracting females at high doses like male pheromones, and attracting males at low doses like female pheromones (Zhang et al. 2008a). In *Tenebrio molitor*, the attractiveness of male pheromones is dose-dependent; at low concentrations, they attract females, but at high concentrations, they become repellent to females (August 1971).

In addition to carboxylic acids, the human skin volatilome of sebum and sweat consists of aldehydes, alkanes, fatty alcohols, ketones, benzenes and derivatives, alkenes, and menthane monoterpenoids (Mitra et al. 2022; Zhang et al. 2022; Zou and Yang 2022; Irga et al. 2024). Low boiling aldehydes such as octanal, nonanal, and decanal, as well as hexadecanol and octadecanol, can also be detected in both colostrum and areolar (Montgomery’s) gland secretions (Doucet et al. 2009). These aldehydes, alkanols, and ketones might be human pheromone components because they have been proven to be components of pheromones or interspecific chemosignals in insects, spiders, birds, rabbits, goats, and rodents, indicating that they possess the chemical properties of pheromones (Hagelin et al. 2003; Schaal et al. 2003; Zhang et al. 2008a; Xiao et al. 2010; Zhang et al. 2010, 2013; El-Sayed 2014; Inagaki et al. 2014; Murata et al. 2014; Zhang and Zhang 2014).

### 3. A Pheromone May Be A Blend of Multiple Components

A pheromone is usually a simple mixture of multiple chemicals (Silverstein et al. 1966, 1967). In mammals, for example, dominant male mice have several preputial gland-secreted volatile compounds that are increased compared to submissive opponents (Novotny et al. 1990; Fang et al. 2016). The additive effects of all these compounds (not one less) increase sexual attractiveness to females (Fang et al. 2016). Two urine-derived pheromone components (2-(sec-butyl)-4,5-dihydrothiazole and dehydro-exo-brevicomin) together promote male-male aggression, induce the estrous cycle in female mice (the Whitten effect), and attract females (Jemiolo et al. 1985, 1986; Novotny et al. 1985).In birds, there are multiple-component male pheromones and interspecific chemosignals (Zhang et al. 2010; Zhang and Zhang 2014). In the spider species *Pholcus beijingensis*, a female pheromone is composed of two volatile components (a blend of hexadecyl acetate and farnesyl acetate) from spider webs, which attract males (Xiao et al. 2009). Thus, we cannot ignore the possibility of multi-component human pheromones, especially a blend of a few fatty acids (Doty 2014; Wyatt 2015).

### 4. The Effects of Pheromones Are Often Dose-Dependent, Especially Those Related to Emotion

In mammals, stress-related pheromones are usually existing compounds with changes in their amounts. For example, in pronghorns (*Antilocapra americana*), 2-pyrrolidinone emitted from rump glands is a putative alerting pheromone (Wood 2002). In the anal glands of small mustelids, sulfur-containing compounds can act to defend against predators by exhaling large amounts of secretions (Brinck et al. 1983). However, low-dose releases or scent marks can be used for intraspecific chemical communication or eavesdropped by prey to assess predation risk (Brinck et al. 1983; Zhang et al. 2003, 2005). In dominant male mice versus subordinate male opponents, urine farnesene levels and sexual attractiveness increase after dominance-submission is established through chronic male-male aggression (Novotny et al. 1990; Fang et al. 2016). Long-term exposure to low doses of cat urine odor (stressor) instead promotes male mice to produce more male pheromones, such as the two farneses that attract females and repel males, rather than inhibiting them (Jemiolo et al. 1991, 1992; Zhang et al. 2008e).

The so-called rat alarm pheromone (2-heptanone) is actually a volatile metabolite of urine origin that is released by a male in an increased way to cause panic in another male rat (Alarm pheromone may evoke panic responses in both sexes, not just the same sex. However, female subjects were not used in the study) (Gutiérrez-García et al. 2007). In fact, 2-heptanone is a male pheromone that attracts females and mediates female mate choice in rats (Zhang et al. 2008b; Zhang and Zhang 2014; Zhang et al. 2019a). This suggests that the same pheromone component may have opposite effects on males and females. For example, in mice, two preputial gland-secreted farnesenes are male pheromones that attract females but repel males (Jemiolo et al. 1991, 1992).

In humans, fear induces higher doses of underarm sweat, releasing more volatile molecules (Lübke et al. 2017). Since humans have emotion-related pheromones, volatile compounds associated with anxiety, fear and stress in human sweat may be some of the existing ingredients, but their amounts will change (Wysocki and Preti 2004; Albrecht et al. 2011; de Groot et al. 2012, 2020; de Groot and Smeets 2017; Wunder et al. 2023). For humans, the increased release of urinary 2-heptanone may be interpreted as an alarm signal indicating imminent danger, similar to its role in other species such as rats, however, human sweat (body odor) does not contain 2-heptanone (Gutiérrez-García et al. 2015). People who are frightened and nervous tend to sweat excessively (hyperhidrosis) and emit a distinctive smell, which may be explained by the increased emanation of existing sweat compounds such as fatty acids (de Groot et al. 2012, 2015, 2020; de Groot and Smeets 2017).

In particular, the effects of predator odor on mice as a typical aversive stimulus are dose-dependent. Low doses of predator odor promote sexual behavior in mice, while high doses inhibit sexual behavior and lead to increased glucocorticoids (Zhang et al. 2003, 2008e; Liu et al. 2017). This further demonstrates that an increase or decrease in the dose of the same chemical component, such as human sweat fatty acids, might cause an opposite change in its effect.

In addition, there is a quantitative differentiation of sex pheromones between geographic populations or strains of the same species (Zhang et al. 2007, 2021, 2023; Zhang and Zhang 2014). For example, in brown rats (Rattus norvegicus), the male pheromone content of populations in cold zones is significantly reduced, and the function of male pheromones is also weakened (Zhang et al. 2021, 2023). Similarly, people of different races exhibit significant differences in odor strength and the functional strength of olfactory communication (Martin et al. 2010).

### 5. Personal Olfactory Information in Human Sweat

Each person emits a unique scent, the most obvious example of which is that dogs can use these scents to identify individuals, track different people, and even distinguish between the odors of identical twins (Kalmus 1955; Penn et al. 2007; Woidtke et al. 2018; Wilson et al. 2022).

The relative standard deviation (RSD = SD/mean) of odor compounds measured by GC-MS can reflect the variability of volatile compositions between individuals, where unusually high RSDs indicate the components’ individuality, as RSDs between duplicate GC-MS experiments are typically <10 (Zhang et al. 2003a, 2005, 2007, 2008d, 2014). In this study, the RSDs of several main fatty acids, such as tetradecanoic acid, pentadecanoic acid, hexadecenoic acid, oleic acid, and octadecanoic acid, were all greater than 30%, indicating that these fatty acids might contribute to coding the olfactory information about the individuality of human sweat odor.

Genetic makeup has a decisive influence on human odor (Wedekind et al. 1995;Thornhill et al. 2003; Preti and Leyden 2010). A functional abcc11 allele is essential for human axillary odor production and related to Caucasians and Africans possessing a strong axillary odor and Asians having only a faint acidic odor (Martin et al. 2010; Preti and Leyden 2010). Some studies illustrated that major histocompatibility complex genes (MHC or HLA) are correlated with body scent attractiveness, mating preference, and likely body odor components as in mice (Wedekind et al. 1995; Ober et al. 1997; Singer et al. 1997;Thornhill et al. 2003; Penn et al. 2007). We also observed that the sweat volatile components differed between Han Chinese and the Chinese of Xinjiang ethnic minorities close to Central Asian countries (unpublished data). It is worth further investigation of how the volatile components of sweat, especially squalene and fatty acids, precisely encode individual odor information and how they relate to genes.

### 6. How Sweat Compounds Affect Menstrual Synchrony and Suppression?

Of the three classical reproductive phenomena in mice (Bruce effect, Whitten effect, and Lee–Boot effect), the latter two can be compared to menstrual synchrony and suppression in women (Bruce 1956; Whitten 1956; Lee and Boot 1959; McClintock 1971; Stern and McClintock 1998).

For the Whitten effect in female mice, it has been demonstrated that male urine-borne pheromones (e.g., the blend of 2-(sec-butyl)-4,5-dihydrothiazole and dehydro-exo-brevicomin, metabolites in urine; or the blend of *E*,*E*-α-farnesene and *E*-β-farnesene secreted from preputial glands) act as male stimuli to induce synchronous estrus in female mice (Whitten 1956; Jemiolo et al. 1985, 1986). For the McClintock effect in women, some women who live in dormitories collectively release more body odor, concentrated in a small, relatively enclosed environment, leading to the accumulation of large amounts of squalene and carboxylic fatty acids in the air. This might be similar to long-term exposure to male body odor, as men release more of these sweat compounds.

It has also been reported that skin ketones, fatty acids, and salivary cortisol increase during menstruation (Fujii et al. 2023). In rhesus monkeys, some fatty acids from vaginal discharges may act as female pheromones to affect male behavior (Curtis et al. 1971; Michael et al. 1971; Bonsall et al. 1980). In women, vaginal excretions contain various fatty acids and lactic acid associated with menstruation (Michael et al. 1974; Bauman et al. 1982). Lactic acid may also have the chemical properties of pheromones, as it acts as an interspecific chemical signal to attract mosquitoes over distances (Acree et al. 1968). Therefore, in addition to the fatty acids in sweat, fatty acids and lactic acid released by the vagina might also play a similar role in olfactory stimulation among roommates in girls’ dormitories.

For the Lee-Boot effect in female mice, which involves the suppression or prolongation of estrous cycles in mature female mice housed in same-sex groups, an essential urine component, 2,5-dimethylpyrazine (possibly an adrenal-dependent compound that may increase in overcrowded females), plays a key role in the urine chemosignals that suppress reproduction through olfaction (Novotny et al. 1986; Ma et al. 1998). In female mice, urine 2-heptanone, trans-5-hepten-2-one, trans-4-hepten-2-one, pentyl acetate, and cis-2-penten-1-yl acetate are also adrenal-dependent but more dependent on estrogen. These compounds vary with the estrous cycle, showing a significant increase on estrus day and a decrease on diestrus days, indicative of potential female pheromones (Schwende et al. 1984; Novotny et al. 1986). We recently found that these five compounds also decreased in overcrowded female mice, consistent with reproductive suppression (Lee and Boot 1959; Schwende et al. 1984; unpublished data). Therefore, further refinement is needed in the study of the pheromones causing the Lee-Boot effect in mice. Nonetheless, the role of 2,5-dimethylpyrazine in the Lee-Boot effect remains well-established (Novotny et al. 1986; Ma et al. 1998).

In fact, 2,5-dimethylpyrazine is also a major component in the urine of ferrets (rodent predators) and acts as an aversive odor and consequent stressor for mice (Zhang et al. 2007; Beny and Kimchi 2016). In women, oral volatile sulfur compounds (halitosis-related malodor) increase during the menstrual and the period blood, such as a metallic or sweaty smell (Huggins and Preti 1981; Dasharathy et al. 2012). Therefore, in addition to the smell from sweat, unpleasant odors from other sources during menstruation might offer a stimulus to suppress reproduction in the McClintock effect.

Of course, it may not be entirely appropriate to compare the Whitten effect and the Lee-Boot effect in mice to the McClintock effect in women. Even in rats, whose phylogenetic relationship with mice is much closer than that of humans, the Bruce effect and the Whitten effect do not occur as they do in mice, and the Lee-Boot effect is not as pronounced in rats as it is in mice (Sharp and LaRegina 1998). It seems that the synchronization, shortening, or delay of the estrous cycle in female rats is closer to the McClintock effect in women, but the key chemical components of the female rat odor at work have not been verified (McClintock,1971, 1978,1984; Stern and McClintock 1998; Zhang et al. 2008; Zhang and Zhang 2014). Anyhow, the relationship between reproductive phenomena and pheromones or odor in mice and rats may encourage us to consider the role of human odor signals and crossmodal olfactory cues to induce the McClintock effect (Spence 2021).

### 7. How Humans Receive Pheromones

The vomeronasal organ (VNO), the sensory organ that initiates the Bruce, Whitten, and Lee-Boot effects in mice, does not exist in humans (Grammer et al. 2005; Brennan and Zufall 2006). The human VNO has epithelia with sensory neurons but does not respond to inhaled volatile pheromones because it has no connections with the central nervous system, the VNO receptor genes are nonfunctional pseudogenes, and the accessory olfactory bulb is present only in the fetus (Grammer et al. 2005).

However, humans rely on the main olfactory system to distinguish between males and females by the smell of sweat (Zhou and Chen 2008). Pheromone receptor genes have been found in the main olfactory mucosa in humans (Rodriguez et al. 2000; Rodriguez and Mombaerts 2002; Grammer et al. 2005). It has been demonstrated that the main olfactory system can also detect pheromones in mice (Wang et al. 1999).

Birds do not have vomeronasal organs but can respond to pheromones via the main olfactory system (Keverne et al. 1999; Hagelin et al. 2003; Brennan and Zufall 2006; Brennan 2010; Zhang et al. 2010, 2013). Even the pheromone androstenone mediates behavioral responses in domestic pigs without passing through the vomeronasal organ (Dorries et al. 1997). These examples illustrate that the vomeronasal organ is not necessary for pheromone detection or pheromone-mediated behavioral responses, even in mammals.

In fact, our human sense of smell is so important and acute that it shapes our social interactions in ways we do not consciously realize, for which, odors enter the brain through the frontal cortex, allowing us to distinguish among different people, such as men and women, and trigger reproduction, emotions, and memories (Glausiusz 2008; Spence 2021). We can subconsciously use smell to assess a person’s sexual attractiveness and emotional state (Glausiusz 2008). In humans, about 300 active olfactory receptor genes are devoted to detecting thousands of different volatile molecules with a molecular weight of less than 300 Da through a large family of olfactory receptors with diverse protein sequences (Sowndhararajan and Kim 2016). Electroneurophysiological tests have revealed that the sense of smell plays an important role in altering cognition, mood, and social behavior (Sowndhararajan and Kim 2016).

## 8. Olfactory Learning Is Important for Pheromone Perception

Social and physical environments can effectively stimulate the human olfactory system to improve olfactory sensitivity (Oleszkiewicz et al. 2021). Conscious focus on and short-term exposure to selected odors may increase olfactory sensitivity (Hummel et al. 2009; Oleszkiewicz et al. 2021). Smell training can improve the sense of smell and alleviate the depression associated with smell loss in humans (von Bothmer 2006; Hummel et al. 2009). This suggests that the odor characteristics of the environment in which people live, as well as the experience and memory of odors, have a significant impact on the sensitivity and perception of smell.

In mice, a major urinary protein (MUP) pheromone, darcin, acts as an innate male pheromone for female attraction and can also condition urine-borne volatile components (learned pheromones) to attract females (Roberts et al. 2010, 2012). In natural populations, social interaction between males and females provides females with learning opportunities for volatile pheromones through associations with MUPs. For example, if female rats are kept in same-sex groups from weaning to puberty, they will lose sexual attraction to male urine odor and the volatile male pheromone 2-heptanone, but they can recover this attraction with activated brain areas (i.e., ventromedial hypothalamus, VMH) that regulate female gender recognition and sexual behavior through learning by associating male urine odor or 2-heptanone with the MUP13 pheromone (Zhang et al. 2019b). The perception of volatile pheromones by the vomeronasal system is also acquired, not just innate (Keverne et al. 1999; Brennan and Zufall 2006; Brennan 2010; Zhang et al. 2019b).

The ability to smell androstenone is genetically determined, and some people who cannot detect the odor of androstenone can learn to smell it through repeated exposures (Wysocki and Beauchamp 1984). In rats, however, females can learn olfactory preferences through repeated exposure to 2-heptanone but do not activate the VMH without associative learning with MUP13 (Zhang et al. 2019b). These studies illustrate the complexity of social learning and its importance for pheromone functions in rodents, which may also apply to all mammals, including humans.

